# Parallel Factor ChIP Provides Essential Internal Control for Quantitative Differential ChIP-Seq

**DOI:** 10.1101/182261

**Authors:** Michael J Guertin, Amy E Cullen, Florian Markowetz, Andrew N Holding

## Abstract

A key challenge in quantitative ChIP-seq is the normalisation of data in the presence of genome-wide changes in occupancy. Analysis-based normalisation methods were developed for transcriptomic data and these are dependent on the underlying assumption that total transcription does not change between conditions. For genome-wide changes in transcription factor binding, these assumptions do not hold true. The challenges in normalisation are confounded by experimental variability during sample preparation, processing, and recovery.

We present a novel normalisation strategy utilising an internal standard of unchanged peaks for reference. Our method can be readily applied to monitor genome-wide changes by ChIP-seq that are otherwise lost or misrepresented through analytical normalisation. We compare our approach to normalisation by total read depth and two alternative methods that utilise external experimental controls to study transcription factor binding. We successfully resolve the key challenges in quantitative ChIP-seq analysis and demonstrate its application by monitoring the loss of Estrogen Receptor-alpha (ER) binding upon fulvestrant treatment, ER binding in response to estrodiol, ER mediated change in H4K12 acetylation and profiling ER binding in Patient-Derived Xenographs. This is supported by an adaptable pipeline to normalise and quantify differential transcription factor binding genome-wide and generate metrics for differential binding at individual sites.

GRAPHICAL ABSTRACT

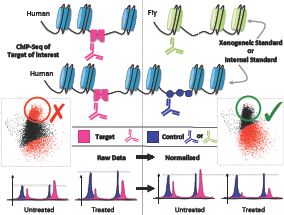

## INTRODUCTION

ChIP combined with high-throughput sequencing (ChIP-seq) quantifies the relative binding intensity of protein/DNA interactions genome-wide for a single condition(1, 2, 3). However, comparing relative intensities of binding between samples and between conditions is an ongoing challenge(4, 5, 6, 7, 8). Conventionally, correcting for sample-to-sample variability between conditions occurs at the analysis stage(9, 10, 11, 12), but these methods assume that experimental variables remain constant between datasets and assume comparable genomic binding of the protein between conditions. In practice, different efficiencies in nuclear extraction, DNA shearing and immunoprecipitation present potential points within a typical ChIP-seq protocol(13) to introduce experimental variation and error(14). Analytical normalisation methods exist to control for variability between samples of the same condition(14, 15), but these methods cannot account for experimental variation between conditions(7). In order to approximate normalisation between conditions the field has exploited a deficiency in ChIP-seq. In short, the total read depth is used as a normalization factor because the vast majority of ChIP-seq reads are outside of true transcription factor binding sites (8, 9). Nonetheless, this approach does not control for any of the aforementioned causes of experimental variability and differences in DNA recovery can be interpreted as differential binding. Previous studies have aimed to resolve these challenges when analysing genome-wide changes through the use of external spike-in controls(4, 5). These methods rely on xenogeneic chromatin (i.e. from a second organism) and either a second species-specific antibody(5), or the cross-reactivity of a single antibody to the factor of interest(4) in both organisms.

Here we present a method, termed parallel-factor ChIP, that utilises a second antibody (anti-CTCF) to provide an internal control. The process of utilising a second antibody against the target chromatin avoids the need of a xenogeneic spike-in and controls for more experimental variables than previous methods. In contrast to spike-in methods, this approach controls for cell lysis conditions, immunoprecipitation efficiency, and sonication fragment size. Moreover, parallel-factor ChIP is not dependent upon accurate quantification of spike-in chromatin. We present this method alongside the application of two xenogeneic methods for the analysis of the fold-change in TF binding between two conditions. Further, we have developed an adaptable pipeline to apply these strategies and provide a highly reliable quantitative analysis of differential binding sites utilising established statistical software packages.

### Estrogen Receptor-alpha as a Model Transcription Factor

Nuclear hormone receptors are a super-family of ligand-activated transcription factors (TF). Many of the molecular mechanisms underlying well-characterised robust and rapidly inducible transcriptional responses, such as estrogen signaling, are shared among other systems. Therefore, we use the transcriptional response to estrogen treatment as a model system to study transcription factor binding. Moreover, many of the aforementioned normalization challenges are exacerbated in the case of ligand inducible TFs (7). For our development and comparison of methods, we monitored ER binding upon treatment with fulvestrant(16). Accurate analysis of the ER binding is of key interest as 70% of all breast cancer tumours are classified as ER+(17). Fulvestrant is a targeted therapeutic to prevent the growth of ER+ tumours(18, 19). The mode of action for fulvestrant is to bind to the ER as an antagonist, which results in recruitment of a different set of cofactors compared to the native ligand estra-2-diol. The fulvestrant-specific cofactors promote degradation of the ER(20, 21) via the ubiquitination pathway and the proteasome(22). The family of compounds to which fulvestrant belongs is called Selective Estrogen Receptor Degraders or Downregulators (SERDs). Cellular loss of ER protein results in compromised ER binding genome-wide and is thus an ideal model for the development of novel quantitative ChIP-seq normalisation methods.

## MATERIALS AND METHODS

### Experimental Design

For experiments containing xenogeneic spike-in material, we generated four replicates for both the control and fulvestrant treatment, a total of 8 samples for the Drosophila spike-in and 8 ChIP-seq samples for the murine chromatin spike-in. For the CTCF parallel-factor ChIP experiments, 3 replicates were prepared for the parallel ER-CTCF pull-down for both control and treatment, giving a total of 6 samples. A single replicate of the CTCF-only pull-down was prepared for both control and treatment conditions.

### Cell Culture

All experimental conditions were conducted in the MCF-7 (Human, ATCC) cell line. Spike-in standards were generated using HC11 (Mouse, ATCC) and S2 (Drosophila, ATCC) cells. MCF-7 were authenticated using STR DNA profiling.

For each individual ChIP pull-down, 4 × 10^7^ MCF-7 cells were cultured asynchronously, as previously described (29), across two 15 cm diameter plates in DMEM (Dulbecco’s Modified Eagle’s Medium, Glibco) with 10% FBS, Glutamine and Penicillin/Streptomycin (Glibco). Incubators were set to 37 °C and to provide a humidified atmosphere with 5% CO_2_.

The cells were treated with either fulvestrant or estradiol (E2) (final concentration 100 nM, Sigma-Aldrich). Prior to E2 treatment, cells were washed with PBS and grown for 4 days in phenol red-free media supplemented with charcoal-stripped FBS. Media was changed daily. The cells were then incubated for the appropriate time period: 48 hours fulvestrant, 2 hours for the effect of E2 on H4K12ac or 45 minutes for ER activation. The cells were washed with ice cold PBS twice and then fixed by incubating with 15 mL per plate of 1% formaldehyde in unsupplemented clear media for 10 minutes. The reaction was stopped by the addition of 1.5 mL of 2.5 M glycine and the plates were washed twice with ice cold PBS. Cells were released from each plate using a cell lifter and 1 mL of PBS with protease inhibitors (PI) into a 1.5 mL microcentrifuge tube. The cells were centrifuged at 8000 rpm for 3 minutes at 4 °C and the supernatant removed. The process was repeated for a second wash in 1 mL PBS+PI and the PBS removed before storing at −80 ^°^C.

S2 Cells were grown in T175 flask with Schneiders Drosophila Medium + 10% FBS at 27 °C. Cells were released by agitation and transferred to a 50 mL Falcon tube. The cells were then pelleted at 1300 rpm for 3 minutes. The media was removed and the cells resuspended in 7.5 mL PBS. In a fume hood, cells were cross-linked by the addition of 7.5mL 2% formaldehyde in unsupplemented clear media. The reaction was stopped with 3 mL of 1M glycine at 10 minutes. The suspension of cells was then centrifuged at 2000 × g for 5 minutes. The cells were then washed twice with 1.5 mL PBS+PI before the PBS+PI was removed and the cells stored at −80 °C.

Untreated HC11 were prepared following the same procedure as MCF-7.

### Chromatin Immunoprecipitation (ChIP)

ChIP was performed as previously reported for cell lines(13) and tissue(29) with the modifications listed below.

For the *D. melanogaster* chromatin spike-in experiment (sequencing data: SLX-8047), *D. melanogaster* and *H. sapiens* samples were prepared separately following the reported protocol until completion of the sonication step. Next, the MCF-7 (experimental) chromatin was combined with the S2 derived chromatin (control) in a ratio of 10:1. Magnetic protein A beads were prepared identically for both the target antibody (100μg, ER, SC-543, lot K0113, Santa Cruz) and the control antibody (10μL, H2Av, 39715, lot 1341001). The washed beads were then combined in a ratio of 1:4 for pull-down. For the *M. musculus* chromatin spike-in experiment (sequencing data: SLX-12998), *M. musculus* and *H. sapiens* cells were prepared separately following the aforementioned protocol until after sonication. Next, we combined the chromatin from the experimental samples (4 × 10^7^ MCF-7 cells) with that from a single plate of HC11 cells (2 × 10^6^ cells). The protocol was continued unmodified using only the ER antibody and protein A beads.

For experiments containing the CTCF antibody control (sequencing data: SLX-14229, SLX-14438, SLX-15090, SLX-15091 &SLX-15439): 100*μ*L magnetic protein G beads were prepared separately for both antibodies, CTCF (10*μ*L, 3418 XP, Cell Signaling) and ER (100*μ*g, SC-543, lots F1716, F0316 and H1216, Santa Cruz) or H4K12ac (100*μ*g, 07-595, Lot: 2884543 Millipore). The beads were then combined 1:1 giving 200*μ*L of beads. The only exceptions were the two CTCF controls (one with and one without treatment) where no ER beads were added. These samples were used to generate a CTCF consensus peak set.

### Library Prep

ChIP and input DNA were processed using the Thruplex Library DNA-seq Kit (Rubicon) according to the manufacturer’s protocol.

### Sequencing

Sequencing was carried out by the CRUK Cambridge Institute Genomics Core Facility using a HiSeq 4000, 50bp single end reads.

### Alignment

Previously, Egan *et al.* (5) aligned the reads to the genomes of the two species separately for the generation of correction factors. We developed our protocol around the alignment to a single combined reference genome, either Drosophila-Human (DmHs) or Mouse-Human (MmHs). The reference genomes were generated from Hg19 and Mm9 or Dm3. We used BowTie2 (version 2.3.2) to align the FASTQ format reads. This resolves and simplifies the challenge of ambiguous alignments between the two genomes. Reads were removed from blacklisted regions (http://mitra.stanford.edu/kundaje/akundaje/release/blacklists/).

### Peak Calling

We used MACS2 (version 2.1.1, default parameters) to call peaks against the combined genome. An example with input data is provided within the Brundle Example repository in Git Hub.

Motif analysis was performed using Homer (v4.9) to provide confidence in peak sets; ER and CTCF control showed a strong enrichment of the full CTCF motif (p-value ≈0). Pairwise IDR (irreproducible discovery rate) analysis of all samples confirmed reproducibility and is summarised in Supplemental Figure S3, S14C and S16C. QC reports are summarised in Table S3.

### qPCR Validation of peaks

Loss or gain of ER binding at known ER binding sites near RARAα, NRIP1 and XBP1 were confirmed by ChIP-qPCR (Figure S4) and changes in H4K12ac was monitored at GREB1, CXCL12 and XBP1. Primers were as previously reported(23, 24, 25, 26). Fold enrichment was calculated against a control region of the genome, proximal to TFF1, known to not bind by ER and to be free of H4K12ac marks from our own ChIP-seq data. The enrichment values were normalised to an input control. The primer sequences for the ER unbound control genomic region were as previously reported(24).

### Bioinformatic Analysis

The bioinformatic analysis was implemented using R (version 3.3.2) with a modified version of DiffBind (version 2.5.6, available from the AndrewHolding/BrundleDevelopment repository on GitHub) and DESeq2 (version 1.14.1). These modifications have been included for the next release of DiffBind from Bioconductor.

### Gene Set Enrichment Analysis

Gene set enrichment analysis of the ER peaks that responded to fulvestrant treatment (FDR=0.01) as established by the parallel-factor ER-CTCF ChIP were submitted to GREAT(27) for analysis. These gave an enriched estrogenic signal (Table S1 and S2).

### Data Availability

All sequence data utilised for this study is available from the Gene Expression Omnibus (GSE102882, GSE107749 and GSE110824).

UCSC Genome browser sessions for the data analysis can be found in the ReadMe.md file uploaded to the AndrewHolding/Brundle R-Package repository on GitHub.

### Pipeline and R packages

An R package containing the functions used for the analysis can be installed directly from CRAN or via AndrewHolding/Brundle on GitHub using the install_github found in the Devtools package.

An R package containing two sets (one internal and one spike-in control) of test data provided as aligned reads, peak files and samples sheets can be installed from AndrewHolding/BrundleData on GitHub.

The complete set of scripts for the preprocessing pipeline is provided to support the implementation of future analysis with Brundle in the preprocessing folder of AndrewHolding/Brundle_Example GitHub. All the contents of the Brundle_Example repository are also packaged in a Docker container for easy use. Instructions on downloading and running the container are available in the ReadMe.md file.

## RESULTS

### Analytical normalisation methods highlight the need for experimental quantitative ChIP-seq controls

Three data-based normalisation strategies are commonly used to normalise ChIP-seq binding between conditions: Reads Per Million (RPM) reads in peaks, RPM total reads, and RPM aligned reads. We applied these methods to each of our ER ChIP-seq datasets to highlight their deficiencies. Despite the presence of spike-in chromatin, these analyses only considered reads that align to the *H. sapiens* genome. CTCF binding sites were excluded from the analysis of parallel-factor ChIP-seq data. We present the analysis of the xenogeneic spike-in and human/mouse cross-reacting ER antibody below, but analysis of all datasets gave consistent results and exhibited a strong decrease in ER binding upon fulvestrant treatment.

We first plotted the average ER peak intensity, as determined by raw counts and three counts-based normalisation methods, by the change in ER intensity upon fulvestrant treatment (Figure S1). In properly normalised MA plots, the unchanged peaks between conditions are distributed with a log-fold difference centered on zero with increasing variance as the peak intensity decreases. However, the distribution of data points in the raw counts MA plot show that this distribution is shifted up to a y-value of ~1 (Figure S1A). We hypothesised that these are true ER binding sites that do not change upon fulvestrant treatment or false-positive peaks. In both cases, the apparent increase in binding would therefore be an artefact of the data processing. As expected, the apparent fold-change for the increase in ER binding was most pronounced when the data was normalised with respect to total number of reads in peaks (Figure S1B) because this method is reliant on the majority of binding events between the two experimental conditions remaining constant. Other common normalisation methods that have been applied to ChIP-seq data, such as quantile normalisation(32, 33), would result in a similar systematic error in the final data. More appropriate methods that correct for total library size, such as Reads Per Million total reads, showed little improvement for our datasets over the raw number of reads counts in peaks (Figure S1C). Each normalisation strategy erroneously implies an increase in ER binding to the chromatin at a large number of sites after 48 hours of treatment.

### Comparison of existing methods

To confirm that the normalisation effects we observed were typical of the commonly used tools for ChIP-seq analysis, we compared results from ChIPComp(28), DiffBind(29), DeSEQ2(31) and EdgeR(30). In a recent comparative analysis, ChIPComp and DiffBind were the only two methods recommended for analysis of narrow peak protein/DNA binding data(12). We therefore compared the results from these two pipelines with EdgeR and DeSeq2, which are routinely applied to ChIP-seq data. The data showed (Figure S2) that ChIPComp, EdgeR and DeSEQ2 detect a large number of significantly unregulated ER binding sites. DiffBind outperformed these methods using total aligned reads for correction. However, Figure S1C highlights the limitations of using total aligned reads.

### Internal and spike-in normalisation controls

#### Normalisation using D. melanogaster chromatin and species-specific antibody for H2Av

To overcome the challenges of normalising ChIP-seq data, Egan *et* al.(5) combined the extract with xenogeneic chromatin and a second antibody that is specific to the spike-in organism’s chromatin. This controls for the efficiency of the immunoprecipitation if the same ratio of target to control chromatin is achieved between samples. This work reported that a reduction in H3K27me3 in response to inhibition of the EZH2 methyltransferase cannot be detected by standard normalisation techniques. Instead, the study demonstrated genomic H3K27me3 reduction by including *D. melanogaster* (Dm) derived chromatin and a Dm-specific histone variant H2Av antibody as a spike-in control for normalisation. However, this method fails to control for variation in sonication fragment length distributions or innaccuracies in quantifying chromatin concentration.

The challenge in analysing the genome-wide reduction in H3K27 methylation by ChIP-seq shares many similarities to quantifying changes in ER binding after fulvestrant treatment.

In particular, both result in a global unidirectional change in chromatin occupancy due to the specific loss of the target molecule.

We applied this method of normalisation to fulvestrant-depleted ER samples using xenogeneic *D. melanogaster* chromatin and an H2Av antibody. Figure 1A shows a similar distribution to Figure S1C, including the off-centre putative unchanged ER binding events (Figure 1A, within red triangle) as highlighted in Figure S1A. Overlaying the peak information from the *D. melanogaster* peaks indicated that they overlapped along the same y-axis value (Figure 1B) as the ER binding events (Figure 1A) that are presumptively unchanged or false positive peaks. We then applied a linear fit to Dm log2(fold-change) values for each binding site. The coefficients generated from the linear regression were then used to adjust the log2 (fold-change) of all data points (Figure 1C). The normalisation of the data resulted in a reduced number of increased ER binding events at 48 hours. The remaining loci of increased binding resulted from the higher variation at lower intensities.

**Figure 1.**
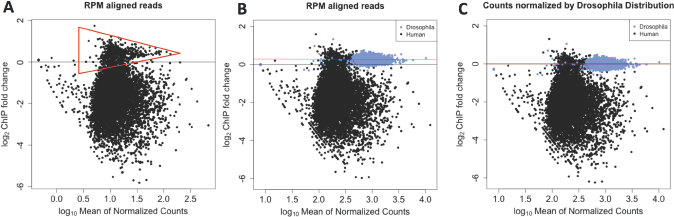
MA plots showing ER binding before and after treatment with fulvestrant including matched Dm H2Av spike-in control. (A) Reads corrected to total aligned reads showed the same off-centre peak density as observed in Figure 1. Putative unchanged ER binding sites are within the red triangle. (B) Overlaying the MA plot combining the changes in chromatin binding of Hs ER (black) and Dm H2Av (blue). Dm peaks overlay the off-centre peak density. (C) Utilising the Dm H2Av binding events as a ground truth for 0-fold change, a linear fit to the log-fold change is generated and the fit is applied to adjust the Hs ER binding events.

#### Normalisation utilising ER antibody cross-reactivity and spike-in murine chromatin

A challenge with using *D. melanogaster* spike-in chromatin as a reference standard for *H. sapiens* ChIP-seq experiments is that both antibody and chromatin must be precisely and accurately quantified. This is technically challenging because cross-linking efficiency, the fragment size and the protein concentration of *H. sapiens* chromatin may not be constant between experimental conditions. In an attempt to reduce the number of variables that can result in experimental error, we developed a similar method to that of Bonhoure *et* al(6). Their study utilised the cross-reactivity of a Pol II antibody against Hs control chromatin and sample chromatin from *Mus musculus* (Mm). The ER antibody utilised in this study is known to cross-react with both Hs and Mm ER homologues. We therefore expected that the inclusion of Mm chromatin would provide a series of control data points that would remain constant between conditions. Unexpectedly, we found that Mm genomic ER peaks were greatly increased after treatment with fulvestrant (Figure S5A). We compared the level of Hs and Mm reads between samples and found the ratios to be consistent (Figure S6), which precludes poor sample balancing as the cause of the results presented in Figure S5.

These results highlight a problem with using a constant antibody and a xenogeneic source of chromatin for normalisation. Despite constant levels of mouse ER, as the spike-in cell line was not treated with fulvestrant, we observe an apparent change in ER binding. We propose that the ER antibody has lower affinity for mouse ER, compared to human ER. Therefore, we conclude that the increase in Mm reads from ER binding sites results from a reduction in competition with human ER for the same antibody, because fulvestrant is degrading human ER. These challenges are likely to be less of a concern when applying this method to a more conserved target and this explains why there has been previous success in applying this strategy to the analysis of histones(5) and RNA Polymerase(6).

#### Normalisation using a second control antibody to provide an internal control

A key reason for utilising the cross-reactivity of antibodies between organisms was to reduce the number of sources for experimental variation. For the same reason, we developed the use of a second antibody as an experimental control to normalise the signal. The advantages of using a second antibody over a spike-in control is that the target:control antibody ratio can be maintained for all samples by producing a single stock solution. For concurrent experiments, a single stock of antibody-bound beads can be prepared and used for all samples with minimal variation. For this control to be effective, it is critical to identify a DNA-binding protein whose genomic distribution and intensities are not affected by the treatment. For the analysis of ER binding, we chose CTCF as our control antibody. While CTCF is affected by compounds that target ER, the effects of these changes have been documented at only a small fraction of the total number of sites(34), a result that was subsequently replicated in our own analysis (Figure 2, Figure S9b and Figure S8).

**Figure 2.**
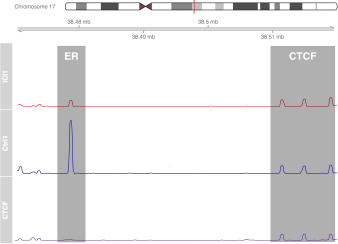
CTCF peak height remains constant while ER peaks change upon treatment with fulvestrant. As the binding of CTCF at the 3 control peaks (right) will remain constant in all 3 conditions (see also Figure 7B), the data is scaled to CTCF peak height. After 100 nM fulvestrant treatment for 48 hours, ER binding (left) shows a reduction in binding at the RARA gene (red) when compared to control (blue). The CTCF peaks can be confirmed against a CTCF only ChIP-seq experiment (red).

We separated the ER and CTCF binding events and plotted them separately on an MA plot (Figure S7A and B). As previously shown for Dm spike-in control, we applied a global fit to the log2-fold change between the two conditions, thereby correcting the bias in fold-change between conditions in ER binding (Figure S7C). Taken together, we show that performing a parallel ChIP-seq experiment with an unrelated and relatively unchanged factor is an alternative and complementary method to account for extreme genomic changes in factor occupancy.

### Pipeline and Quantitative Analysis

#### H2Av and CTCF provide a set of unchanged reference peaks for normalisation

For a parallel-factor ChIP to be effective as an internal control, the majority of the binding sites for the control factor must not change between the two conditions. We identified control CTCF peaks from a conventional CTCF ChIP-seq experiment that did not include ER antibody. Since the signal at CTCF-proximal ER binding sites may change upon fulvestrant treatment due to the overlapping signal from ER peaks, we excluded all CTCF ChIP-seq peaks that are within 500bp of previously identified ER binding sites from MCF-7 cells. Comparison of the two control datasets (Figure S9) displayed a lower variance and a lower maximum fold-change for H2Av compared to the CTCF control binding regions. In contrast, the CTCF dataset provides a much greater number of data points for normalisation as a result of relative size of the human and Drosophila genomes. None of the H2Av sites in the Drosophila genome or CTCF sites used for normalisation showed a significant change in occupancy.

#### Normalisation implementation using DESeq2 and Size Factors

DESeq2 was initially developed for the analysis of RNA-seq data(31) to provide a method to quantify significant differences in gene expression between two samples by modelling gene counts data with a negative binomial distribution. Given the similarities in ChIP-seq and RNA-seq, primarily that they are both based on the same high throughput sequencing technologies, DESeq2 has been successfully adapted to ChIP-seq analysis to establish differential intensity analysis of histone modifications.

DESeq2 is designed for an RNA-seq library where total transcription is assumed to not change between conditions and ~100% of counts are signal (in contrast, the ChIP-seq signal is often contributed by fewer than 5% of reads). As expected, the default DESeq2 *estimateSizeFactors()* parameter calculated from a ChIP-seq counts table distorted the average change in ER signal because the assumption of constant total binding between conditions is not met (Figure S10A). In the dual antibody Dm spike-in experiment, the Dm H2Av peaks should be constant. We manually used the read counts in these H2Av control peaks as a size factors parameter estimate for correcting ER binding intensities (Figure S10B). We processed the CTCF internal control data in the same manner, using the counts with CTCF peaks to adjust the size factors parameter. We normalised the data using the counts within CTCF peaks to estimate the DESeq2 size factors (Figure S11B).

#### Integration with DiffBind using Corrected Size Factors

DiffBind(29) is an established R package to provide a pipeline to quantitatively measure differential binding from ChIP-seq data. DiffBind has been applied to a variety of ChIP-seq studies; recent examples include the epigenomic landscapes of retinal rods and codes(35), the interaction of MDM2 polycomb repressor complex 2(36), and establishing an environmental stress response network in *Arabidopsis*(37). In a comparative study of ChIP-seq analysis tools, DiffBind reliably outperformed other methods(12) and is the preferred strategy for analysis of ChIP-seq experiments with multiple replicates. For these reasons, we chose DiffBind to underpin our analytical methodology and as a key benchmark to improve upon. A key feature of DiffBind is that, to calculate size factors, it utilises the total library size from the sequence data provided in a sample sheet (e.g. BAM files) rather than the *estimateSizeFactors* function provided by DESeq2. Nonetheless, while improved, the analysis of the raw data by DiffBind is incomplete with the putative unchanging peaks showing a greater than 0 log-fold change (Figure S12). To address this shortcoming, we modified the DiffBind package to directly calculate the *sizeFactor* parameter from a counts matrix of control peaks, in our case either H2Av or CTCF peaks (Figure S12B).

### Establishing a normalisation coefficient by linear regression of control peak counts

DESeq2 generates the size factor estimates through the summation of all reads within the peaks, resulting in a bias to the peaks with the largest read count. We therefore hypothesised that we could improve normalisation by calculating the sample bias through the application of linear regression. We plot the read count in each CTCF peak of one condition against the other (Figure 3) and then apply a linear model to the data. Our normalisation coefficient is defined as the constant by which we need to scale the count data for each CTCF peak from the treated samples to correct this systematic bias (and thereby setting the gradient of the linear fit equal to 1). This normalisation coefficient is then applied in the same manner to ER count data and then reinserted into the DiffBind object for analysis.

**Figure 3.**
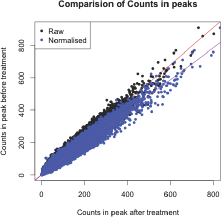
Comparison of mean counts in CTCF peaks before and after treatment. If the samples have no systematic bias before and after treatment then the linear fit would be expected to have a gradient of 1. Here, we establish that the gradient is < 1, implying a systematic bias between samples. The read counts in the treated samples peaks are corrected (blue), removing the bias, and resulting in a new gradient of 1.

We compared normalisation by total library size, CTCF control peak-derived size factors, and linear regression to our sample data. Our linear regression method provided higher sensitivity, as 10.7% more sites were detected as differentially bound (FDR < 0.05) compared to normalisation by library size alone (Figure 4).

**Figure 4.**
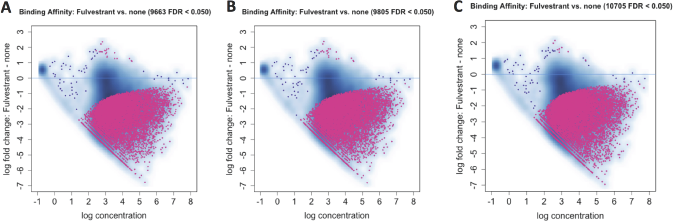
Comparison of DiffBind results before and after our two methods of normalisation. (A) Normalisation to Library Size. (B) Applying the corrected size factors from our DESeq2 pipeline generated from CTCF internal control. (C) Applying correction using linear regression of CTCF peaks between conditions to normalise data. The result is a 10.7% increase in the number of loci detected as significantly changed ER binding.

### Normalisation factors are consistent over a wide range in number of control binding sites

In order to determine if parallel-factor ChIP normalisation could be used with factors that are not pervasively bound throughout the genome like CTCF, we recalculated the normalisation coefficient by sub-sampling from 100 to 1% of the CTCF peaks. The variability of the result was then modelled by re-sampling each analysis 100 times (Figure 5). When sampling only 1% of sites at random 50% cases resulted in an error of less than 0.5% and the maximum error was still within 2% of the expected value. This analysis indicates that parallel factor ChIP is robust and that the number of control peaks can vary over two orders of magnitude and not substantially affect the normalisation factor.

**Figure 5.**
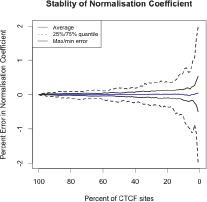
Stability of CTCF derived normalisation coefficient. Stability of the CTCF derived normalisation coefficient was analysed by sub-sampling CTCF peaks before undertaking the calculation (between 1-100% of total sites) at random. This analysis was repeated 100 times to model the variability of the result.

### Normalisation of samples with minimal binding condition

In the absence of E2, ER binding to DNA is nearly undetectable by ChIP-seq. The minimal level of TF binding in the initial condition could present a challenge to normalisation. To confirm if parallel-factor ChIP was suitable for application to conditions with a very low level of initial binding, we applied our pipeline to the analysis of ER binding in E2-free conditions and 45 minutes after stimulation with 100 nM E2. The data was normalised using our pipeline and we identified 16884 sites of significantly increased binding (FDR = 0.05, Figure S14A). Analysis of normalised read depth at known binding sites near RARα, NRIP1, and XBP1 genes showed an increase in ER binding as expected (Figure S14B). Comparison of conditions show good correlation between replicates (Figure S14C). Motif analysis of the sites displaying significantly increased binding gave strong enrichment for the motifs of the ERE, FOXA1 and GATA3 representing the core ER complex (Figure S14D). Comparison of sites that showed increased ER binding (FDR=0.01, Figure S15) overlapped with a core of 1312 conserved sites across 4 independent studies and > 60% of peaks overlapped with at least 1 other dataset.

### Parallel-Factor ChIP to normalise broad histone modification peaks

Applying parallel-factor ChIP to histone modifications presents an additional challenge because histone modifications occur over broad domains, as opposed to the discrete binding TFs. To demonstrate the application of parallel-factor ChIP-seq to histone marks, we applied our method to H4K12ac in MCF7 cells. ER regulates H4K12 acetylation through the recruitment of BRD4(26). Analysis of the normalised data showed an increase of H4K12ac at 11393 sites and reduction at 4817 sites (Figure S16A), overall resulting in a significant increase (p-value = 5.7 ×10^−12^) of the H4K12ac histone mark as expected. 377 of the individual sites are significant after multiple testing correction (FDR = 0.05). As no genome-wide statistical analysis had previously been undertaken at individual peaks, we cannot compare this result; however, included in those 377 significant sites were GREB1 (FDR = 2.7 ×10^−4^) and XBP1 (FDR = 3.0 ×10^−6^) peaks near their respective TSS, as previously reported(26). Analysis by qPCR of H4K12ac of GREB1, CXCL12 and XBP1 sites (Figure S16B), along with the H4K12ac occupancy profile +/-3000bp of ER Binding (Figure S16D), agreed with a previous report(26). As H4K12ac is commonly associated with transcription(38) and previous work reported that ER recruits BRD4 to increase H4K12 acetylation at active promoters(26), we repeated the analysis focusing on H4K12ac occupancy within +/- 500 bp of ER binding at transcription start sites. Under this more stringent filtering, we identified 497 ER promoter regions with H4K12ac occupancy. Of these sites, 28 regions were found to have significantly increased levels of H4K12ac compared to 5 regions with decreased (FDR = 0.05) occupancy, equating to ~6-fold more sites with increased H4K12ac than had decreased. In comparison, we observed a ~2-fold bias genome-wide.

### Comparison of absolute fold-change from parallel-factor ChIP and xenogeneic spike-in

A small subset of high-intensity low-fold change peaks, i.e. those at the narrow end of the triangle in Figure 1A, were absent in the MA plots of samples generated with the parallel pull-down of CTCF and ER (Figure 1A and S7A). To address if masking of ER binding sites by CTCF has a significant impact on the results of ER parallel-factor ChIP, we re-analysed the data using a consensus set of 10,000 high-confidence ER binding sites (as established by ER-only ChIP). Normalisation was carried out as previously described, either using the Dm chromatin or the CTCF loci. In principle, if both the internal control using CTCF binding events and the use of the spike-in Dm/H2Av control are accurate, the normalised fold-change for each genomic loci between the two data sets should be equal. Plotting the fold-change of normalised results from the two experimental methods (Figure 6) gave a result of near parity between the methods (linear fit of gradient = 0.94) and a correlation of r = 0.77, with a p-value tending to 0).

**Figure 6.**
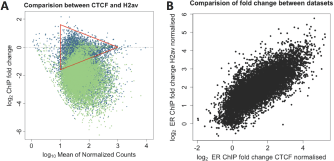
Comparison of normalisation methods using consensus peak set. (A) The analysis for the CTCF normalised (blue) and H2Av normalised (green) dataset using an ER consensus peak set of 10,000 peaks were formatted as an MA plot and overlaid. This recovered the low-fold change higher-intensity peaks that were not visible in Figure S7A and both datasets showed a similar distribution. (B) Comparison of fold-change values for individual ER binding sites between two datasets showed that the inclusion of these sites did not appear to affect the correlation (r = 0.77).

### Cross-normalisation of Single-Factor ChIP to Parallel-Factor ChIP

A potential limitation of parallel-factor ChIP is that CTCF sites may suppress the fold-change measurement of proximal TF binding sites. To address this, we made use of an intrinsic feature of standard ChIP-seq that the method accurately quantifies relative binding intensities within the same pulldown. By quantifying TF binding at sites that are not proximal to CTCF in a parallel-factor ChIP experiment, we can normalise all sites in a TF-only ChIP. To demonstrate cross-normalisation, we used the Hs reads from the HsDm dataset as an example of an ER-only ChIP-seq dataset. We established a set of consensus peaks by matching non-CTCF proximal ER binding sites from our parallel-factor ChIP with ER binding sites in our ER-only experiment. Given that relative binding between sites is intrinsically accurate by normalising the ER consensus site binding in the ER-only experiment to the normalised parallel-factor ChIP, we were able to accurately normalise all sites in the ER-only experiment. As the ER-only data we used contained xenogeneic spike-in controls, We were able to validate the cross-normalisation. Comparison of log-fold change after normalisation using the xenogeneic spike-in and cross-normalisation showed cross-normalisation gave equivalent results to that previously seen: Pearson’s correlation of 0.992, p-value tending to 0 (Figure S13A). Analysis of ER binding events proximal to CTCF after cross-normalisation showed a marginally greater magnitude of mean and maximal fold-change compared to that established by parallel-factor ChIP (Figure S13B). We can therefore ascertain that cross-normalisation provides a robust strategy to establish changes in TF binding; however, in the case of ER binding, the suppressive effect of proximal CTCF binding is minimal.

### Analysis of Patient-Derived Xenografts (PDX) by Parallel-Factor ChIP

To demonstrate the versatility of parallel-factor ChIP-seq, we applied our method to the analysis of five PDX samples. The analysis of PDXs presents similar challenges to that of clinical material. As a consequence of the high levels of sample heterogeneity, the sample preparation and immunoprecipitation steps in the ChIP protocol are significantly more variable than for cell lines. The low amounts and the high value of samples present further challenges by limiting the ability to perform replicate experiemnts and analysis.

Analysis of CTCF binding within the samples acted as a QC step (Figure S17A). PDX02 showed no enrichment at either CTCF or ER binding sites, thereby confirming the result was not due to low-expression of ER in the PDX material. The sample was therefore excluded from further analysis. Clustering of samples by ER binding events gave two clusters with PDX01 and PDX04 displaying the greatest correlation (Figure S17B). A potential reason for the clustering is PDX01 and PDX04 are both derived from PR positive tumours, while PDX05 is derived from a PR negative tumour. The PR status of PDX03 is unknown.

Comparison of normalisation to total read count (RPM) and parallel-factor ChIP showed a large disparity between the two methods at the RARAa, GREB1 and CLIC6 ER binding sites (Figure S17C). Analysis of the variance of the CTCF control peaks proximal to these sites demonstrated Parallel-Factor ChIP-seq was able to stabilise the data (Figure S17D) while normalisation to total read count gave little improvement over the raw data. PDX05 was found to have the lowest levels of ER bound at the sites investigated.

Genome wide profiling of the parallel-factor ChIP-seq PDX data was in agreement with the analysis of individal promoters. CTCF binding was normalised between samples (Figure S18 top) and gave a consistent profile. ER binding genome-wide was then normalised on the basis of the correction established from CTCF binding. Before normalisation all four samples displayed different maximum levels of ER binding. After normalisation PDX01, PDX03 and PDX04 gave similar levels of ER binding, all derived from tumours with an Allred(39) score of 8 (a immunohistochemical score out of 8 estimating the proportion and intensity of ER-staining in tumor cells). In agreement with the analysis of RARAa, GREB1 and CLIC6 ER binding sites, PDX05’s binding profile showed a reduce maximum level of binding. These results is in agreement with the PDX05 being derived from a tumour with Allred score of 5 (Figure S18 bottom).

## DISCUSSION

We have described a normalisation strategy using internal ChIP-seq controls. We applied this technique to normalise TF binding in a model system and patient derived xenograft smaples. Moreover, we developed and implemented a statistical analysis at the level of individual binding sites, which was lacking from previous spike-in methodologies. We demonstrate that a parallel-factor control antibody is a reliable alternative to previously described experimental controls(4, 5).

We showed that an internal parallel-factor control is comparably quantitative to using a second antibody and xenogeneic chromatin as a spike-in control, but there are many advantages to using a second antibody (CTCF) that IPs a protein within the same extract. Primarily, the parallel-factor ChIP controls for the greatest number of steps in the process and gives fewer opportunities for variation being introduced into the sample preparation. In contrast, the addition of xenogeneic chromatin relies on the precision that the concentration of the chromatin of both the experimental samples and the spike-in can be established reliably and must be added to each sample individually. As chromatin is routinely cross-linked for ChIP-seq, the resultant mixture of protein and DNA makes accurate quantification of DNA challenging without purification, which presents another challenge for the use of xenogeneic spike-in methods.

### Limitations

Normalization has over-promised the ability to directly compare different ChIP-seq experimental conditions, an aim that is intrinsically not possible due to the inherent biological and environmental variability between experiments. As a result, inconsistency between ER ChIP-seq in previous datasets is an ongoing challenge(48). While parallel-factor ChIP provides an essential normalisation between conditions, it should be understood that the method cannot control for large-scale biological and environmental factors. Comparisons of our ER binding data with previously published data gave a core 1312 conserved ER binding sites between all datasets (Figure S15). Improving on a previous comparison of similar data sets which gave only 284(48). Overlap of our these data was substantial at >60%; nonetheless, this analysis highlights the need to understand that biological variability is distinct from the technical variability for which parallel-factor ChIP is designed to control. The key challenge our method resolves is providing a value of fold-change from differential analysis that is accurate and comparable between experiments, which has not previously been possible with analytical normalisation(7). Once the fold-change for each peak has been established, then we can undertake direct comparison of fold-change between datasets through the use of consensus peak sets(Figure 6).

The reliability of any experimental control is critical for any normalisation technique. For the parallel-factor ChIP peaks, we undertook triplicate biological replicates. If one was to require the CTCF peak to appear in every replicate, this would result in over 54000 high confidence peaks in our test dataset. Analysis of the stability of the normalisation coefficient showed only a small fraction of this number of sites is needed with less than a 2% maximal error when using only 1% of CTCF peaks (Figure 5). Nonetheless, due to the key role that normalisation plays in the downstream data analysis, the quality of the data obtained should be assessed by a QC pipeline, e.g. ChIPQC(44) and NGS-QC(45).

Importantly, our use of normalisation controls appear resilient to changes in antibody batch. There are genuine concerns in reproducibility of ChIP-seq as a result of batch variation in antibody. We were able to demonstrate strong correlation between the xenogeneic spike-in and parallel-factor controls despite the two experiments being conducted with different lots of ER antibody (see Methods) and at different times. Nonetheless, the initial differential analysis that establishes normalised fold-change should be performed with the same batch and source of antibody.

Parallel-factor ChIP has broad utility in the chromatin and transcription fields. First, we established the ability to normalise signal from samples that have effectively no detectable binding in the initial condition. We exhibited this ability using the extreme example of a nuclear receptor that is nearly entirely unbound in the ligand-free condition. Secondly, we showed that this approach effectively normalizes histone modification ChIP-seq data, which presents a distinct set of challenges(7). We were able to reliably normalise both ER and H4K12ac ChIP-seq signal to the control factor that was immunoprecipitated in parallel (CTCF). Previous studies provided evidence of a global increase in H4K12ac. Through the application of parallel-factor ChIP, we were able to monitor changes in individual regions of H4K12ac genome-wide. In agreement with Nagarajan *et al.*(26), we found average occupancy of H4K12ac increases; however, we showed the increase is coupled with a global redistribution of H4K12ac not previously described. Analytical normalisation would typically suppress measurement of the global increase in H4K12ac, yet the use of parallel-factor ChIP enabled the quantitative analysis of the increase in the H4K12ac histone mark while simultaneously providing evidence of the redistribution of H4K12ac histone occupancy. This example exemplifies the power of the internal controls provided by parallel-factor ChIP. Without these controls, we would have been unable to reconcile our more detailed analysis with the results presented by Nagarajan *et al.*

### Experimental normalisation is essential and complementary to analytical normalisation

Normalisation at the analysis stage has developed considerably since early ChIP-seq experiments; recent examples include ChIPComp(28), csaw(11) and HMCan-diff(15). In contrast to analytical normalisation, the development of experimental sample controls is more limited(4, 5, 6, 8). Experimental normalisation, including parallel-factor controls, remain necessary as analytical normalisation of pull-down efficiency is only possible between replicates of the same explicit condition(14, 15). Without experimental controls to provide a reference, any systematic bias between conditions will remain indistinguishable from biological signal.

### ER response to fulvestrant

The only previous ChIP-seq study of the effects of fulvestrant on ER binding(47) identified 10205 ER binding sites in the control condition. The ER binding was compared to tamoxifen (8855 peaks) and fulvestrant (4285 peaks) treatments, and concluded the presence of ligand-specific binding. This result has since been disputed in the context of the tamoxifen treatment(49). The majority of the tamoxifen-specific peaks were reassigned as ER peaks by Hurtado *et al.* and, of the remaining tamoxifen-specific sites, only 7 were found in both studies and therefore not reproducible. Our analysis of ER binding identified 13745 sites in the control condition under the more stringent requirements. After normalisation, we found no evidence that fulvestrant induced ligand-specific binding at 48h after treatment. Given a single replicate, it is not possible to establish a statistical test of binding at each site from the Welboren *et al.* data set. Our analysis found 10705 (FDR <0.05) differentially bound sites, which is substantially more than previously identified. Gene Set Enrichment Analysis with GREAT(27) confirmed consistency with the literature as there was significant enrichment for the ER pathways for both the MSigDB pathway and perturbation datasets.

### Importance of experimental normalisation

Normalisation has played a key role in these analyses as, before normalisation, our analysis found sites that would be considered to have significantly increased ER binding on fulvestrant treatment. Further, as we repeated the experiment with two different normalisation techniques, we can confidently state that in the context of asynchronous MCF7 cells, fulvestrant does not result in any significantly increased binding after 48 hours of treatment.

We have shown, as parallel-factor ChIP-seq utilises internal standards, our protocol can be applied to the analysis of tumour samples, PDXs and other clinical material. Consistent sample preparation is a key challenge in clinical sample studies and by controlling for variation in cell lysis, immunoprecipitation, and sonication efficiency, parallel-factor ChIP allows for the deconvolution of biological signal from variability in sample preparation in a way that is not possible with spike-in normalisation methods. As implemented here, one could monitor if individuals who are heterozygous for DNA binding proteins have absolute reduced binding or if the absolute levels of TF binding increase during disease progression.

### Integration with existing methods

Most importantly, we have developed the analysis tools to integrate the normalisation strategies described into well-established quantitative ChIP-seq analysis methods(31). By providing an open and reproducible pipeline, we permit others the ability to accurately normalise transcription factor binding. We expect future studies of transcription factors that undergo rapid and genome-wide changes will find the methods we present essential to accurately characterise biological effects. Our analysis tools, combined with the benefits and relative simplicity of parallel-factor ChIP to normalise ChIP-seq data, have provided a fundamental resource for quantitative transcription factor analysis.

## FUNDING

We would like to acknowledge the support of the University of Cambridge, Cancer Research UK and Hutchison Whampoa Limited.

Parts of this work were funded by CRUK core grant [grant numbers C14303/A17197, A19274] to FM; Breast Cancer Now Award [grant number 2012NovPR042] to FM; CRUK Travel Award [grant number C60571/A24631] to AH; and a Thomas Jefferson Fellowship to AH.

## ACKNOWLEDGEMENTS

MJG and ANH conceived the normalisation strategies, designed the experiments, undertook the data analysis, and wrote the manuscript. ANH and AEC undertook the ChIP-seq experiments. ANH developed the R data and analysis packages. FM reviewed and contributed to the initial proposal and provided feedback to the project.

We would like to acknowledge the contribution from the CRUK Genomics and Bioinformatic core facilities in supporting this work.

We are grateful to Rory Stark for supplying a modified version of his DiffBind Package and to Ashley Sawle and Federico Giorgi for their ideas and support in strategies for alignments to multiple genomes.

We would like to thank Caldas Lab at CRUK CI for providing PDX material to support our method development.

## Conflict of interest statement

None declared.

